# Ultrasound Imaging of Gene Expression in Mammalian Cells

**DOI:** 10.1101/580647

**Authors:** Arash Farhadi, Gabrielle H. Ho, Daniel P. Sawyer, Raymond W. Bourdeau, Mikhail G. Shapiro

## Abstract

The study of cellular processes occurring inside intact organisms and the development of cell-based diagnostic and therapeutic agents requires methods to visualize cellular functions such as gene expression in deep tissues. Ultrasound is a widely used biomedical technology enabling deep-tissue imaging with high spatial and temporal resolution. However, no genetically encoded molecular reporters are available to connect ultrasound contrast to gene expression in mammalian cells. To address this limitation, we introduce the first mammalian acoustic reporter genes. Starting with an eleven-gene polycistronic gene cluster derived from bacteria, we engineered a eukaryotic genetic program whose introduction into mammalian cells results in the expression of a unique class of intracellular air-filled protein nanostructures called gas vesicles. The scattering of ultrasound by these nanostructures allows mammalian cells to be visualized at volumetric densities below 0.5%, enables the monitoring of dynamic circuit-driven gene expression, and permits high-resolution imaging of gene expression in living animals. These mammalian acoustic reporter genes enable previously impossible approaches to monitoring the location, viability and function of mammalian cells *in vivo*.

## INTRODUCTION

The study of cellular function within the context of intact living organisms is a grand challenge in biological research and synthetic biology (*1*). Addressing this challenge requires imaging tools to visualize specific cells in tissues ranging from the developing brain to tumors, and to monitor gene- and cell-based therapeutic agents *in vivo* (*2*). However, most common methods for imaging cellular processes such as gene expression rely on fluorescent or luminescent proteins, which have limited performance in intact animals due to the poor penetration of light in biological tissue (*3, 4*). On the other hand, ultrasound easily penetrates most tissues, enabling deep non-invasive imaging with excellent spatial and temporal resolution (∼100 µm and ∼1 ms, respectively) (*2, 5*). These capabilities, along with its safety, portability and low cost, have made ultrasound one of the most widely used technologies in biomedicine. Despite these advantages, to date ultrasound has played a relatively small role in cellular imaging due to the lack of appropriate genetically encoded reporters.

Recently, the first biomolecular contrast agents for ultrasound were introduced based on gas vesicles, air-filled protein nanostructures which evolved in certain waterborne bacteria and archaea to provide cellular buoyancy (*6, 7*). Gas vesicles comprise a 2 nm-thick protein shell enclosing a gas compartment with dimensions on the order of 100 nm (**Fig. 1**). The acoustic impedance mismatch between their gas interior and surrounding aqueous media allows gas vesicles to strongly scatter sound waves and thereby serve as ultrasound contrast agents (*8-12*). In their native organisms, gas vesicles are encoded by clusters of 8-14 genes, including one or two structural proteins that make up the bulk of the gas vesicle shell, and several other essential genes encoding assembly factors or minor shell constituents.

**Fig 1.**
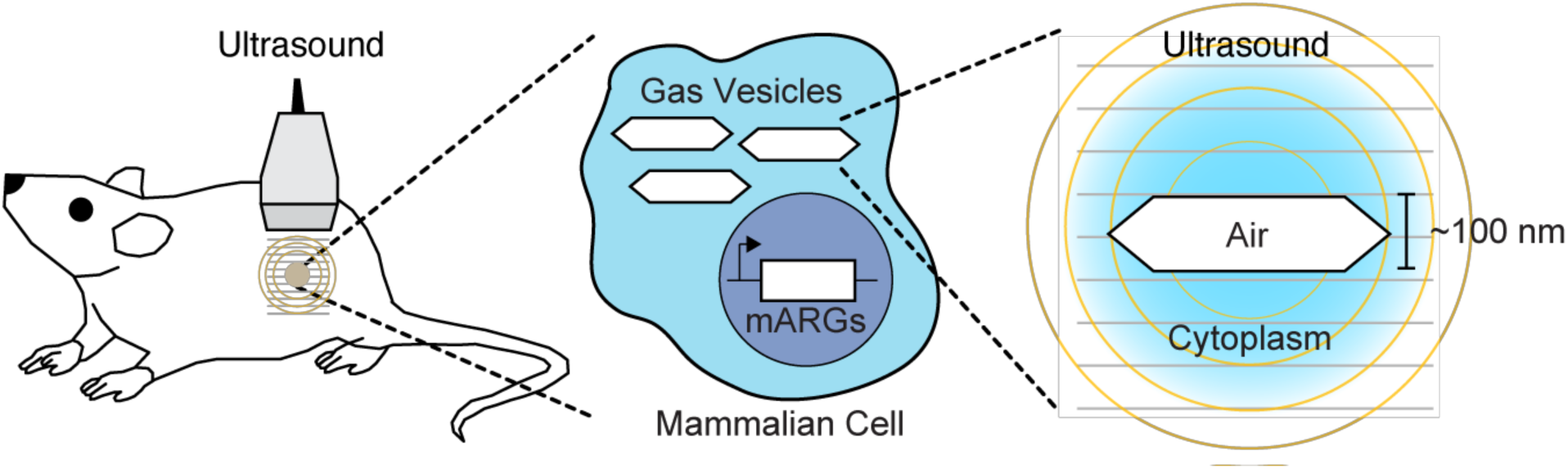
Illustration of mammalian acoustic reporter genes. Mammalian acoustic reporter genes (mARGs) encode a set of proteins whose expression results in the formation of cytoplasmic gas vesicles – air-filled protein nanostructures which scatter ultrasound waves and thereby produce contrast in ultrasound images. This technology allows ultrasound to image mammalian gene expression non-invasively in intact animals.

The use of gas vesicles as reporter genes requires the heterologous expression of their cognate multi-gene operon in a new cellular host, ensuring proper transcription and translation of each gene, functional folding of each corresponding protein and appropriate stoichiometry and co-localization of the constituents for gas vesicle assembly. Recently, a major genetic engineering effort succeeded in expressing gas vesicles as the first acoustic reporter genes (ARGs) in commensal bacteria, allowing their imaging in the mouse gastrointestinal tract (*13*). However, ARGs for mammalian cells have not been developed and represent an even greater synthetic biology challenge. This challenge arises, first, due to the differences in transcription and translation between prokaryotes and eukaryotes. For example, whereas bacterial operons can have multiple genes arranged as sequential open reading frames driven by a shared promoter, this does not normally happen in eukaryotes. Second, the products of individual genes transferred from bacteria to mammalian cells may not fold or function properly due to differences in the cellular environment and availability of chaperones or other cofactors (*14*). Third, for self-assembling multi-gene structures such as gas vesicles, it may be difficult to obtain the correct stoichiometry to enable functional assembly. Finally, while the proteins whose genes share an operon may be spatially co-localized in bacteria due to local translation (*15, 16*), the products of genes expressed separately in mammalian cells may have less efficient co-localization in the relatively vast cytoplasm. To our knowledge, no genetic operon larger than 6 genes has been moved between these domains of life (*17*).

Here, we describe the results of a major effort to express ARGs in mammalian cells and enable ultrasound imaging of mammalian gene expression (**Fig. 1**). Using a stochastic multi-gene screening technique, we identified a set of 9 genes originating from *Bacillus megaterium* that are necessary and sufficient to produce gas vesicles in cultured mammalian cells, as demonstrated with electron microscopy. The engineering of these genes into synthetic operons using viral co-translational elements and the tuning of gene stoichiometry resulted in robust gas vesicle expression in stable mammalian cell lines. Using an ultrasound imaging paradigm maximizing the sensitivity of gas vesicle detection, we were able to image the expression of these ARGs in mammalian cells at volumetric densities below 0.5%, track the dynamics of a representative genetic circuit, and map the precise spatial pattern of gene expression *in vivo* in a murine tumor xenograft.

## RESULTS

### Identifying mammalian acoustic reporter genes

To identify a set of genes capable of encoding gas vesicle assembly in mammalian cells, we used a stochastic gene mixture assay. We synthesized individual gas vesicle genes from three different microbial species using codons optimized for human expression, cloned each gene into a unique monocistronic plasmid and transiently co-transfected mixtures of the genes from each species into HEK293T cells using polyethylenimine (PEI) complexes (**Fig. 2A**). This assay uses two stochastic events to sample a broad range of gene stoichiometries and expression levels in a single experiment. First, each PEI complex incorporates a random number of copies of each plasmid from the mixture, drawn from a distribution centered on that plasmid’s relative concentration. Second, variable numbers of PEI complexes reach a given cell nucleus. This is expected to result in transfected cells receiving a range of random stoichiometries and gene doses in each co-transfection experiment (**Supplementary Fig. 1**).

**Fig 2.**
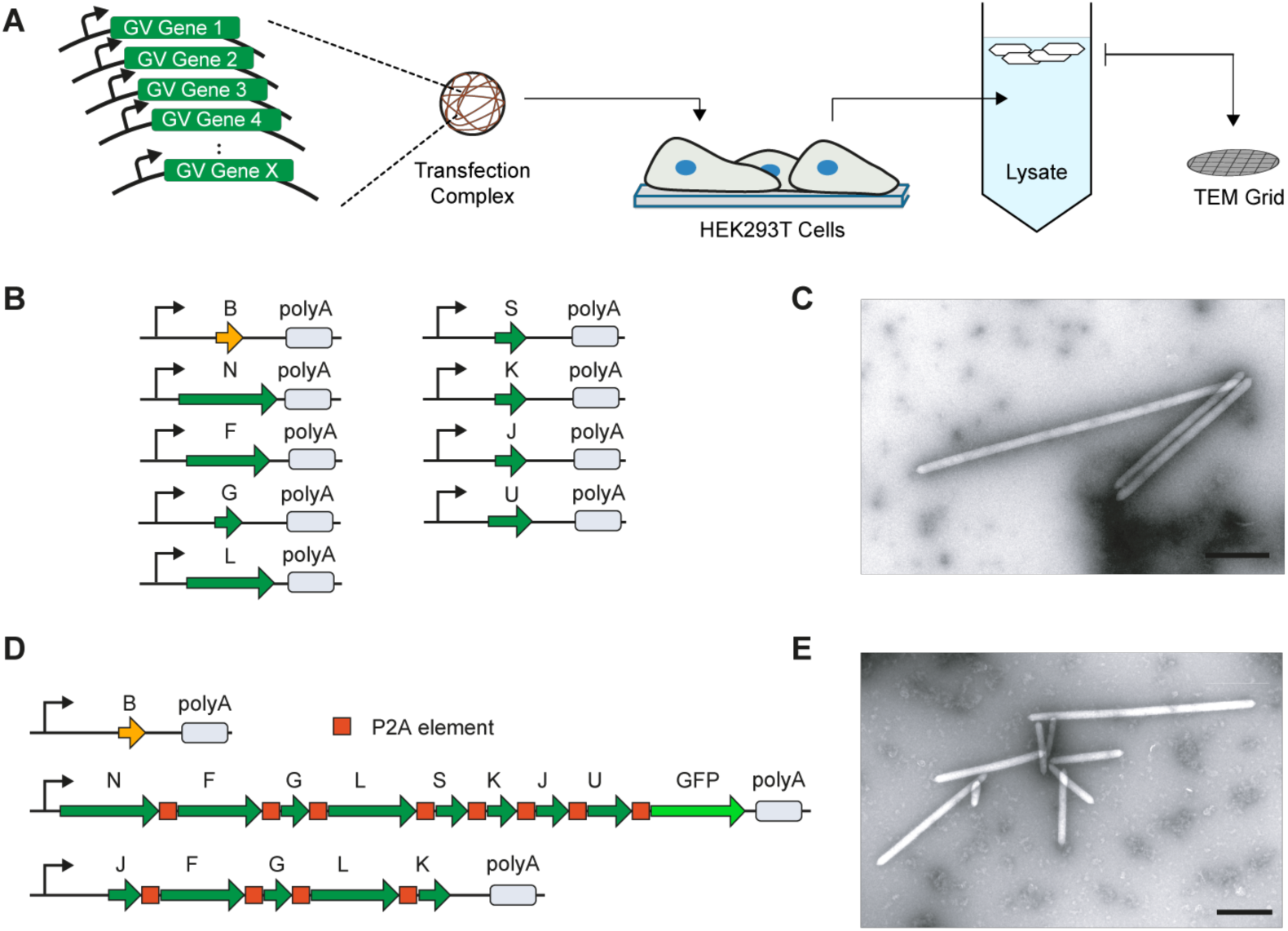
Engineering of mammalian acoustic reporter genes. (**A**) Schematic of the transient co-transfection assay used to identify combinations of genes capable of producing gas vesicles in mammalian cells and the buoyancy-enriched TEM method used to detect gas vesicle formation. (**B**) Schematic of nine genes from *B. megaterium* capable of encoding gas vesicle expression in mammalian cells. Each gene was codon optimized and cloned on an individual plasmid containing a CMV promoter upstream and SV40 polyadenylation element downstream of the gene. (**C**) Representative TEM image of purified gas vesicles expressed in HEK293T cells. (**D**) Synthetic mammalian operon containing 3 gas vesicle gene cassettes that together encode the expression of gas vesicles. The first plasmid encodes the primary structural protein GvpB; the second plasmid encodes all the remaining assembly factor proteins (GvpNFGLSKJU) connected using P2A elements; the third plasmid is a booster cassette encoding GvpJFGLK connected using P2A elements. The three plasmids are each driven by CMV promoters and contain an SV40 polyadenylation element downstream of the last gene. (**E**) Representative TEM image of purified gas vesicles when HEK293T cells are transfected with the constructs in (D). All scale bars represent 500 nm.

After giving the cells 72 hours for protein expression, we gently lysed the cells, and centrifugated the lysate to enrich for buoyant particles, which would include any fully-formed gas vesicles. The top fraction of the centrifugated lysate was then imaged by transmission electron microscopy (TEM). We hypothesized that if gas vesicle expression in HEK293T cells is possible using some combination and ratio of gas vesicle genes, then some cells in the culture would receive this combination, resulting in the presence of gas vesicles in our TEM images.

The co-transfection of the gas vesicle genes from *Halobacterium salinarum* and *Anabaena flos-aquae* in HEK293T cells did not lead to the formation of detectable gas vesicles. However, the co-transfection of 9 gas vesicle-forming genes from *Bacillus megaterium* (**Fig. 2B**) resulted in the production of unmistakable gas vesicles as evidenced by their appearance and geometry in TEM images (**Fig. 2C**). The 9 genes originate from an eleven-gene *B. megaterium* gene cluster previously used to express gas vesicles in *E. coli* (*13, 18*), with the exception of *GvpR* and *GvpT*, which were found to be unnecessary for gas vesicle formation (**Supplementary Fig. 2**).

### Constructing a mammalian operon for gas vesicle expression

Using the 9 genes identified in our stochastic screen, we next set out to construct a polycistronic mammalian operon for consistent assembly of gas vesicles with a compact genetic footprint. To achieve this goal, we joined groups of gas vesicle genes using the viral 2A self-cleavage peptide (*19*) – a sequence encoding 18-25 amino acids that when placed in-frame between two genes causes a ‘ribosomal skip’ on the mRNA, releasing the first protein and proceeding to translate the second protein. The 2A element has a smaller genetic footprint and higher co-expression efficiency than internal ribosome entry sequences, another commonly used polycistronic element. However, the use of 2A elements results in terminal peptide additions to the upstream and downstream proteins, with the C-terminus of the preceding protein receiving a 24-amino acid peptide and the N-terminus of the succeeding protein receiving a proline residue. To determine if the gas vesicle proteins could tolerate these modifications, we added sequences encoding N-terminal prolines and the C-terminal overhangs of the porcine teschovirus-1 2A element (P2A) to each of the 9 *B. megaterium* genes identified in our co-transfection screen (**Supplementary Fig. 3**), and tested the ability of *E. coli* with gene clusters containing these modified genes to form gas vesicles. We found that all genes except *GvpB* tolerated N- and C-terminal P2A modifications (**Supplementary Table 1**).

**Fig 3.**
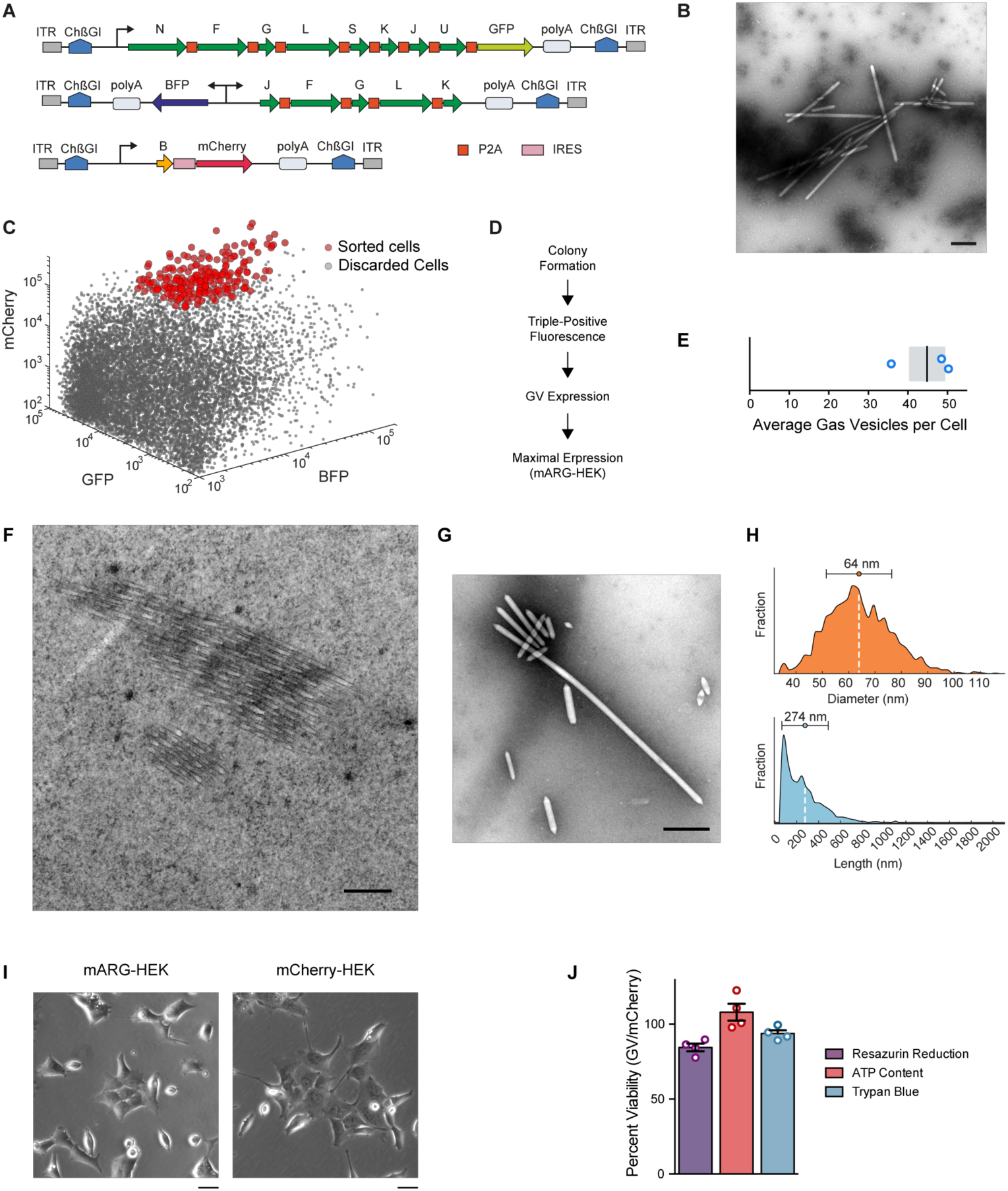
Formation, properties and non-toxicity of gas vesicles in cells with genome-integrated mammalian acoustic reporter genes. (**A**) Schematic of mARG constructs used for genomic integration into cells with the piggyBac transposase system. ITR, inverted terminal repeat; ChβGI, Chicken beta-globin insulator; GFP, Emerald green fluorescent protein; BFP, enhanced blue fluorescent protein 2. (**B**) Representative TEM image of buoyancy-enriched lysate from HEK293-tetON cells transfected with the constructs in (A) and sorted for high expression of all three operons. (**C**) Fluorescence-activated cell sorting of HEK293-tetON cells transfected with the constructs in (A). Red circles denote individual cells selected by sorting to form monoclonal cell lines. (**D**) Selection process for monoclonal cell lines, including assays for viability, fluorescence intensity and gas vesicle yield. (**E**) Number of gas vesicles expressed by monoclonal HEK293-tetON cells after 72 hours of induced expression, as counted in lysates using TEM. Bar represents the mean and the shaded area represents SEM (n=3, each from two technical replicates). (**F**) Representative TEM image of a 60-nm section through an mARG-HEK cell showing an angled slice through two bundles of gas vesicles in the cytosol. (**G**) Representative TEM image of gas vesicles purified from mARG-HEK cells. (**H**) Size distribution of gas vesicles expressed in mARG-HEK cells. The mean and standard deviation of both distributions is illustrated as a circle and with error bars. (n=1828) (**I**) Phase contrast images of mARG-HEK and mCherry-HEK cells 72 hours after induction with 1 µg/mL doxycycline and 5 mM sodium butyrate. (**J**) Cell viability of mARG-HEK cells relative to mCherry-HEK cells after 72 hours of gene expression. Error bars indicate SEM. Scale bars in **B**, **F**, **G** represent 500 nm. Scale bar in **I** represents 20 µm.

Accordingly, we constructed a polycistronic plasmid containing the 8 P2A-tolerant gas vesicle genes connected by P2A sequences, and co-transfected it into HEK293T cells together with a separate plasmid encoding *GvpB*. Unfortunately, this did not result in the production of gas vesicles as assessed by TEM. We hypothesized that one or more of the proteins encoded on the polycistronic plasmid may be expressed at an insufficient level in this genetic arrangement. We tested this hypothesis gene-by-gene by co-transfecting all but one of the 8 gas vesicle genes as monocistronic plasmids, while complementing that one gene solely using our polycistronic 8-gene construct (**Supplementary Fig. 4**). Using TEM to analyze the relative efficiency of gas vesicle formation for each of the 8 genes, we found that *GvpN, GvpS* and *GvpU* supplied in either monocistronic or polycistronic form supported abundant gas vesicle assembly. However, the production of gas vesicles was significantly reduced when *GvpJ, GvpF, GvpG, GvpL* or *GvpK* was supplied from the polycistronic vector. We therefore suspected that these genes represented a bottleneck in gas vesicle formation. To address this limitation, we constructed a polycistronic “booster” plasmid containing these five genes connected by P2A elements. Co-transfection of this plasmid with the full 8-gene polycistronic plasmid and a plasmid encoding *GvpB* (**Fig. 2D**) enabled robust expression of gas vesicles in cells (**Fig. 2E**). We named this set of three genetic constructs mammalian acoustic reporter genes, or mARGs.

**Fig 4.**
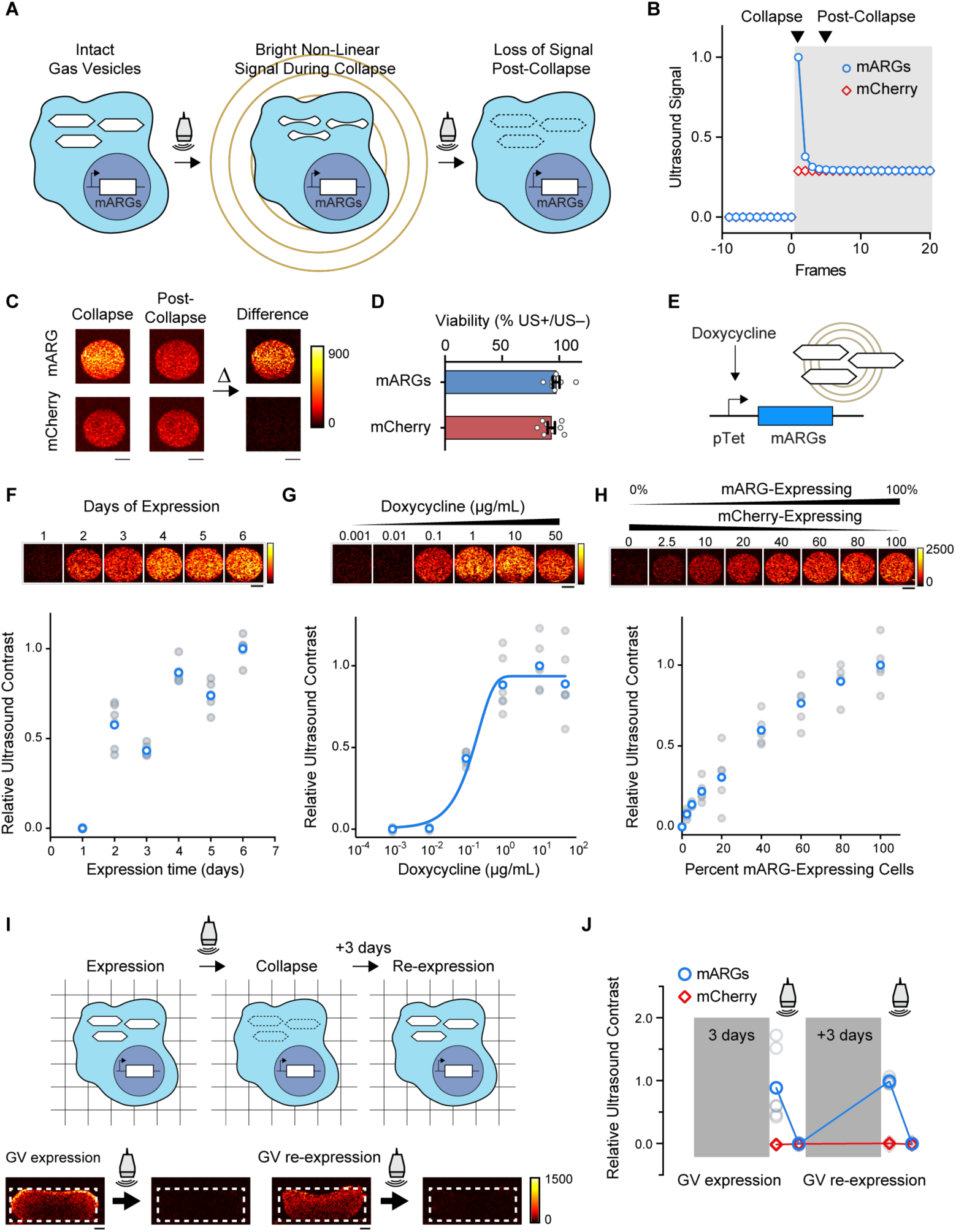
Ultrasound imaging of mammalian gene expression *in vitro*. (**A**) Illustration of the collapse-based ultrasound imaging paradigm used to generate gas vesicle-specific ultrasound contrast from mARG-expressing cells. (**B**) Representative non-linear signal recorded during a step change in the incident acoustic pressure, from 0.27 MPa in the white-shaded region to 1.57 MPa in the grey-shaded region. (**C**) Representative collapse and post-collapse ultrasound images of mARG-HEK and mCherry-HEK cells acquired during this ultrasound imaging paradigm and their difference, indicating gas vesicle-specific contrast. (**D**) Cellular viability after being insonated under 3.2 MPa acoustic pressures, as measured using the MTT assay. (**E**) Schematic of a chemically inducible gene circuit with mARG expression as its output. All three mARG operons in mARG-HEK cells are under the control of the doxycycline-inducible TRE3G promoter (pTet), with expression triggered by incubation with doxycycline. (**F**) Representative ultrasound images and contrast measurements in mARG-HEK cells as a function of time following induction with 1 µg/mL of doxycycline and 5 mM sodium butyrate (n=6, with the darker dots showing the mean). (**G**) Representative ultrasound images and contrast measurements in mARG-HEK cells as a function of doxycycline induction concentrations. Cells were allowed to express gas vesicles for 72 hours in the presence of 5 mM sodium butyrate. (n=6, with the darker dots showing the mean). A sigmoidal function is fitted as a visual guide. (**H**) Representative ultrasound images and contrast measurements in mARG-HEK cells mixed with mCherry-HEK cells in varying proportions. Cells were induced with 1 µg/mL of doxycycline and 5 mM sodium butyrate for 72 hours prior to imaging. (n=5, with the darker dots showing the mean) (**I**) Schematic and representative ultrasound images from mARG-HEK cells in Matrigel re-expressing gas vesicles after acoustic collapse. Cells were induced with 1 µg/mL of doxycycline and 5 mM sodium butyrate for 72 hours before and after 3.2 MPa acoustic insonation. Ultrasound images were acquired after an additional 72 hours in culture following collapse. (**J**) Ultrasound contrast in mARG-HEK and mCherry-HEK cells after initial expression, after collapse, after re-expression and after second collapse. (n=7, with the darker dots showing the mean). GV, gas vesicles. All scale bars represent 1 mm.

### Stable genomic integration of mammalian ARGs

After establishing polycistronic constructs for mammalian gas vesicle assembly, we sought to integrate them into the cellular genome for stable expression. For this purpose, we cloned the constructs into piggyBac integration vectors (*20, 21*) under a doxycycline-inducible TRE3G promoter, with a unique fluorescent protein added as a transfection indicator to each genetic construct (**Fig. 3A**). We co-transfected these plasmids into HEK293-tetON cells together with the piggyBac integrase, and used flow cytometry to sort cells according to their expression level of each fluorescent channel. We found that the cell population combining the strongest expression of each construct produced the largest quantity of gas vesicles (**Fig. 3B, Supplementary Fig. 5, A-D**). To ensure that gas vesicle expression was not unique to HEK293 cells, we performed the same experiment with Chinese hamster ovary cells (CHO-K1), and obtained similar results (**Supplementary Fig. 5, E-G**).

**Fig 5.**
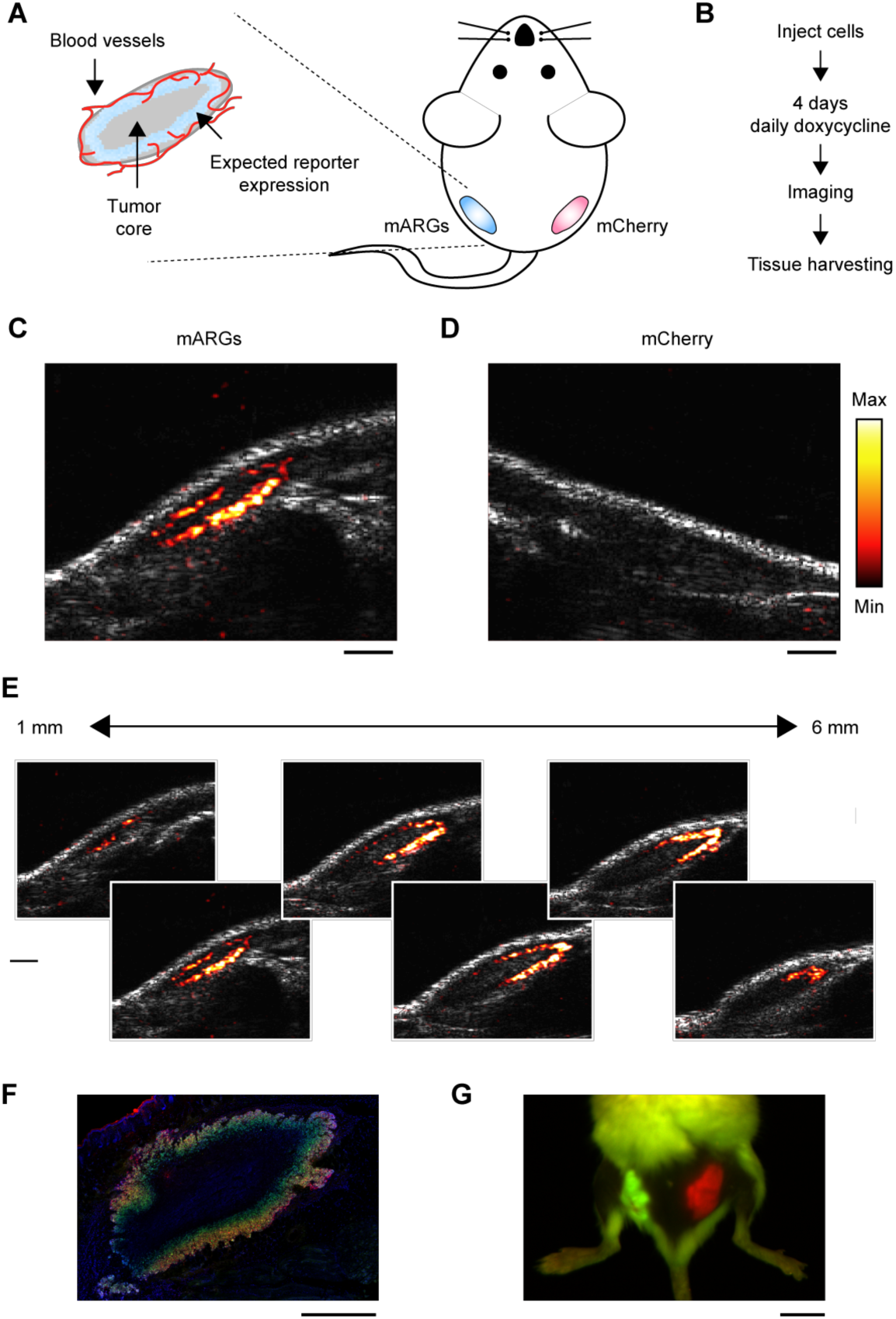
Ultrasound imaging of mammalian gene expression *in vivo*. (**A**) Diagram of a mouse implanted with a subcutaneous tumor model, and the expected spatial pattern of vascularization and doxycycline-induced reporter gene expression. (**B**) Experimental timeline. (**C**) Representative ultrasound image of tumors containing mARG-HEK cells after 4 days of doxycycline administration. mARG-specific contrast shown in the hot colormap is overlaid on an anatomical B-mode image showing the background anatomy. (**D**) Representative ultrasound image of tumors containing mCherry-HEK cells after 4 days of doxycycline administration. (**E**) Ultrasound images of adjacent planes in the mARG-HEK tumor acquired at 1 mm intervals. The minimum and maximum values of color bars in **C-E** are 4000 and 40000 au, respectively. (**F**) Representative fluorescence image of a histological tissue section of an mARG-HEK tumor. Blue color shows the TO-PRO3 nucleus stain, green color shows GFP fluorescence and red color shows mCherry fluorescence. (**G**) Fluorescence image of a mouse implanted with mARG-HEK and mCherry-HEK tumors on the left and right flanks, respectively, after 4 days of expression. Scale bars for are 1 mm for **C-F** and 1 cm for **G**.

To generate a stable monoclonal cell line expressing mARGs for detailed analysis, we sorted individual high-expression HEK293-tetON cells for monoclonal growth (**Fig. 3C**), producing 30 cell lines, which we screened for viability, fluorescence and gas vesicle formation (**Fig. 3D**, **Supplementary Table 2**). The number of gas vesicles per cell was then estimated from TEM images, and a cell line yielding the largest quantity of gas vesicles was selected and named mARG-HEK. When induced for 72 hours with 1 µg/mL of doxycycline and 5 mM sodium butyrate (to prevent epigenetic silencing), this cell line produced on average 45 gas vesicles per cell (**Fig. 3E**). Using thin-section TEM, gas vesicles could clearly be seen in the cytosol of individual mARG-HEK cells (**Fig. 3F**). From TEM images of cell lysates, we measured the average dimensions of gas vesicles produced in this cell line to be 64 ± 12 nm wide (standard deviation, n=1828) and 274 ± 212 nm long (standard deviation, n=1828), with some reaching lengths greater than 1 micron (aspect ratios greater than 30) (**Fig. 3, G-H**). This corresponds to an average gas vesicle volume of 0.605 attoliters. Together, the 45 gas vesicles expressed in an average mARG-HEK cell are expected to occupy just 0.0027% of the cell’s cytosolic volume.

The expression of gas vesicles did not change the gross morphology of mARG-HEK cells (**Fig. 3I**), and was non-toxic as determined by three different assays, including membrane integrity with trypan blue, the relative number of metabolically active cells with CellTiter-Glo, and metabolic activity using resazurin reduction (**Fig. 3J**). For these assays, mARG-HEK cells were compared to an mCherry-expressing control cell line (mCherry-HEK) that was similarly generated using the piggyBac integrase vector in HEK293-tetON cells (**Supplementary Fig. 6 A-B**).

### Ultrasound imaging of mammalian ARG expression

Having established stable gas vesicle expression in a mammalian cell line, we sought to image this expression with ultrasound. Gas vesicles encoded by the *B. megaterium* gene cluster are expected to produce linear ultrasound scattering, emitting the same ultrasound frequency as applied by the transducer with amplitude linearly dependent on the incident pressure (*18*). However, since mammalian cells themselves also produce significant linear scattering, detecting gas vesicles expressed in such cells using linear methods is challenging. To enable more selective imaging of mARG-expression, we took advantage of the ability of gas vesicles to collapse irreversibly above specific ultrasound pressure thresholds (*8, 9, 13, 18*). A switch in the incident ultrasound pressure from below to above such a threshold results in a strong transient signal from the gas vesicles, which decays to a lower level in the next ultrasound frame due to immediate dissolution of their gas contents and the elimination of ultrasound scattering (**Fig. 4, A-B**). Meanwhile, background tissue scattering rises with the increase in incident pressure and remains constant at the new level. Thus, images formed by taking the difference in signal between the collapsing and post-collapse frames reveal specifically the presence of gas vesicles.

We implemented this collapse-based imaging approach using an amplitude modulation pulse sequence (*10*), which we found to provide the best cancellation of non-gas vesicle signals. When hydrogels containing mARG-HEK cells were imaged using this technique at 18 MHz, they were easily distinguishable from mCherry-HEK controls based on their contrast dynamics (**Fig. 4C**). Critically, while this imaging paradigm requires the collapse of gas vesicles inside cells, this does not affect cell viability (**Fig. 4D**).

Reporter genes are often used to track the dynamic activity of natural or synthetic genetic programs (*22, 23*). To test if mARGs can faithfully monitor circuit-driven gene expression, we measured the dynamic ultrasound response of mARG-HEK cells under the control of a doxycycline-inducible promoter (**Fig. 4E**). After induction with 1 µg/mL doxycycline, the cells showed a gradual buildup of ultrasound signal, with clear contrast appearing on day two and increasing over the next 4 days (**Fig. 4F**). These kinetics are similar to those observed with fluorescent indicators (**Supplementary Fig. 7A**). When the gene circuit was driven using a range of input inducer concentrations, the ultrasound contrast produced by cells also followed the expected transfer function of the promoter (**Fig. 4G, Supplementary Fig. 7B**).

Next, we asked how sensitively mARG-expressing cells could be detected in a mixed cell population. To answer this question, we combined mARG-HEK cells with mCherry-HEK cells at varying ratios. We were able to sensitively detect the presence of mARG-expressing cells in these mixtures down to 2.5% of total cells (**Fig. 4H**), corresponding to less than 0.5% volumetric densities, or approximately 4 cells per voxel for voxel dimensions of 100 µm. This sensitivity should be sufficient for a wide variety of biological scenarios.

In many imaging experiments, the output of a gene circuit is read out only once. However, in some cases it may be desirable to track gene expression in a given cell population over time. We therefore tested whether mARG-expressing cells in which the gas vesicles have been collapsed during imaging could re-express these reporters to allow additional imaging at a later timepoint. We embedded mARG-HEK cells in a nutrient-supported three-dimensional hydrogel and imaged them over two sessions. First, the cells were imaged 3 days after induction, showing clear ultrasound contrast. Then the same cells were allowed to re-express gas vesicles over 3 additional days and re-imaged, again showing robust contrast (**Fig. 4, I-J**). The ability of cells to repeatedly express gas vesicles after they are erased with ultrasound will enable longitudinal tracking of cell location and gene expression, and could also be used in pulse-chase experiments visualizing expression or migration dynamics.

### Mammalian ARGs enable ultrasound imaging of gene expression in vivo

Having engineered mammalian cells to stably express gas vesicles and characterized their ability to produce ultrasound contrast *in vitro*, we next tested the ability of mARG expression to be visualized *in vivo* with high spatial resolution. For this purpose, we formed model tumor xenografts in immunocompromised mice by inoculating mARG-HEK cells in Matrigel subcutaneously in their left flanks (**Fig. 5A**). In the same mice, the right flanks were inoculated with mCherry-HEK control cells. We induced reporter gene expression in both tumors for 4 days via systemic injections of doxycycline and sodium butyrate (**Fig. 5B**). We expected these nascent tumors to be mostly vascularized at their perimeter and therefore receive the largest concentrations of doxycycline and oxygen at this location. We therefore hypothesized that this should result in the strongest gene expression at the tumor periphery (**Fig. 5A**). Ultrasound, with its sub-100-µm spatial resolution (at 18 MHz), should be able to discern this gene expression pattern, whereas attaining such resolution would be challenging with optical techniques.

After 4 days of induction, we observed clear ultrasound contrast in the flank inoculated with mARG-HEK cells, which was absent from the contralateral side (**Fig. 5, C-D**). As expected, the pattern observed with ultrasound revealed mARG expression at the perimeter of the tumor, while the core remained dark, and the imaging of adjacent ultrasound planes revealed this pattern of gene expression to persist across the tumor mass (**Fig. 5E, Supplementary Fig. 8**).

The ultrasound-observed spatial distribution of gene expression was consistent with the low vascularity in the tumor core, as observed with Doppler ultrasound (**Supplementary Fig. 9**). The gene expression pattern was confirmed with subsequent histological examination of the tissue using optical microscopy, showing a distinct pattern of peripheral expression (**Fig. 5F, Supplementary Fig. 10**). In comparison, our *in vivo* fluorescence images just showed the presence of signal somewhere in the tissue and not its precise distribution (**Fig. 5G**). These results, which were consistent across 5 animals (**Supplementary Fig. 11**), demonstrate that mARGs enable gene expression imaging *in vivo* and highlight the ability of ultrasound to visualize intricate patterns of gene expression non-invasively with 100-µm spatial resolution.

## DISCUSSION

Our results establish the ability of an engineered genetic construct encoding prokaryote-derived gas vesicles to serve as a mammalian reporter gene for ultrasound, giving this widely used non-invasive imaging modality the ability to monitor cellular location and function inside living organisms. mARGs provide many of the capabilities associated with established genetically encoded optical reporters. This includes the imaging of gene expression over time, the recording of cellular dynamics such as promoter-driven expression activity, and the spatial mapping of gene expression in complex samples. While optical reporter genes mainly provide these capabilities in culture and sectioned or surgically accessed tissues, the acoustic reporter genes described in this work enable gene expression to be visualized non-invasively *in vivo* with 100-µm spatial resolution.

Furthermore, the synthetic biology methods used in this study to transfer the 11-gene polycistronic cluster encoding gas vesicles from prokaryotes to eukaryotes should be instructive for future efforts to import complex genetic machinery from bacteria and archaea into mammalian cells. Our results show that a stochastic expression screen can be used to identify a set of genes that, at some ratio, enables a multi-gene functionality to be recapitulated, that at least 9 genes can be strung together with 2A elements, that polycistronic complementation can be used to identify assembly-limiting genes, and that these limitations can be rectified with a “booster”.

While the mARG constructs described in this work should be immediately useful in a variety of contexts, significant scope exists for further study and optimization to make acoustic reporter genes as widely useful as GFP (*5, 12*). First, while this study used immortalized cell lines and clonal selection to demonstrate gas vesicle formation, future studies should examine the use of mARGs in primary cells. To facilitate such use, it would be helpful to further condense the constructs into a single plasmid. Preliminary experiments show that *gvpB* can be combined with the 8-gene polycistron encoding *gvpN-gvpU* via an internal ribosome entry sequence (**Supplementary Fig. 12**). At 9.5 kb, the total length of the coding sequence included in the three mARG constructs is small enough to fit into a single adenoviral vector. With further optimization to remove the requirement of some genes to be present in two copies, the resulting 7 kb sequence could be small enough for lentiviral delivery.

In this study, we could readily visualize mARG-expression in living cells and animals using a novel ultrasound imaging sequence taking advantage of gas vesicles’ pressure-dependent collapse behavior. This provides impressive sensitivity, with the smallest concentration of mARG-HEK cells imaged in this study corresponding to a 0.5% volume fraction. However, the need to collapse gas vesicles in order to image them limits opportunities for temporal averaging and rapid dynamic monitoring. Prokaryote-derived gas vesicles have been recently engineered at the genetic level to exhibit non-linear acoustic scattering through reversible buckling deformations (*9*), thereby enabling selective imaging without the need for irreversible collapse (*10, 11*). Similar engineering could in the future enable mARGs to be detected non-destructively. Just as the genetic engineering of fluorescent proteins and concurrent advances in optical techniques produced a cornucopia of powerful tools for the study of cells under a microscope, further engineering of acoustic reporter genes and improved acoustic methods for their detection will give rise to new ways to visualize cellular function inside the body.

## MATERIALS AND METHODS

### Chemicals, cell lines and synthesized DNA

All chemicals were purchased form Sigma Aldrich unless otherwise noted. HEK293T and CHO-K1 cell lines were ordered from American Type Culture Collection (ATCC) and HEK293-tetON cells and CHO-tetON cells were purchased form Clontech (Takara Bio). Synthetic DNA was ordered from Twist Bioscience.

### Cloning

Monocistronic plasmids used for transient transfection of HEK293T cells of gas vesicle genes used the pCMVSport backbone. Codon optimized gas vesicle genes were assembled in each plasmid using Gibson assembly. To test the effect of N- and C-terminal P2A modification each *B. megaterium* gas vesicle gene on the pNL29 plasmid (addgene 91696) was individually cloned using standard mutagenesis techniques. To test the N-terminal modification, the CCT codon was inserted following the start codon. To test the C-terminal modification, a linker-P2A sequence (GGAGCGCCAGGTTCCGGG-GCTACTA ACTTCAGCCTCCTTAAACAGGCCGGCGACGTGGAAGAGAATCCTGGC) was inserted upstream of the stop codon for each gene.

The polycistronic plasmid containing gvpN, gvpF, gvpG, gvpL, gvpS, gvpK, gvpJ, gvpU and Emerald GFP (EmGFP) were codon optimized, and synthesized in three fragments. The three fragments were Gibson assembled in the pCMVSport plasmid. The booster plasmid was assembled by multi-fragment Gibson assembly from PCR amplified fragments of the above plasmid.

The piggyBac transposon system (System Biosciences) was used to genomically integrate the mARG cassettes. To clone the mARG cassettes to the piggyBac transposon backbone, the plasmid was first digested using the SpeI and HpaI restriction enzymes and the mARG cassettes were Gibson assembled with the backbone. For doxycycline-inducible expression, the CMV promoter upstream of the gas vesicle genes was replaced with the TRE3G promoter. Internal ribosome entry site (IRES) and mCherry were cloned downstream gvpB as a marker for genomic integration. For the booster plasmid, CMVmin followed by enhanced BFP2 (eBFP2) and a polyadenylation element were cloned in the reverse direction upstream of the TRE3G promoter (creating a bi-directional doxycycline-inducible promoter) and used as a marker for genomic integration. A piggyBac transposon plasmid containing TRE3G and mCherry was Gibson assembled similarly to above.

### Cell culture, transient transfection and TEM analysis

HEK293T and CHO-K1 cells were cultured in DMEM with 10% FBS and penicillin/streptomycin and seeded in a 6-well plate for transfection experiments. When the cells reached 70-80% confluency, 2 µg of total DNA (comprising the indicated mixtures of plasmids) was complexed with 2.58 µg polyethyleneimine (PEI-MAX; Polysciences Inc.) per µg of DNA, added to the cell culture, and incubated for 12-18 hours. Thereafter, the media containing the PEI-DNA complex was changed with fresh media. Cells were allowed to express the recombinant proteins for 72 hours.

To look for gas vesicles, cells cultured in 6-well plates were lysed with 400 µL of Solulyse-M (Genlantis Inc) per well for one hour at 4 °C. The lysate was then transferred to 2 mL tubes, diluted with 800 µL of 10 mM HEPES buffer at pH 8.0 and centrifugated overnight at 300 g and 8 °C. Then, 60 µL of the supernatant was transferred to a fresh tube to be analyzed using transmission electron microscopy (TEM).

From this top fraction, 2 µL of sample was added to Formvar/carbon 200 mesh grids (Ted Pella) that were rendered hydrophilic by glow discharging (Emitek K100X). The samples were then stained with 2% uranyl acetate. The samples were imaged on a FEI Tecnai T12 transmission electron microscope equipped with a Gatan Ultrascan CCD.

To estimate gas vesicle yield and analyze size distribution, the cells were seeded in 6-well plates and gas vesicle expression was induced with 1 µg/mL of doxycycline and 5 mM sodium butyrate for 72 hours. The cells were lysed using Solulyse-M and buoyancy enriched at 300 g at 8 °C overnight. The top fraction of the supernatant was mixed with 2M urea and spotted on Formvar/carbon grids. The TEM grids were washed with water before staining with 2% uranyl acetate. To calculate gas vesicle yield per cell, the total number of gas vesicles per sub-grid on the TEM grid was manually counted and related via lysate volume to the number of source cells. Gas vesicle size distribution was quantified using FIJI.

To visualize gas vesicles inside cells, mARG-HEK cells were seeded in 6-well plates and allowed to express gas vesicles for 72 hours. The cells were fixed in 1.25% glutaraldehyde in PBS, post-fixed in 1% aqueous osmium tetroxide, reduced with ferrocyanide and block-stained in 1% uranyl acetate (all reagents from Electron Microscopy Sciences). The material was then dehydrated through a graded ethanol series and embedded in Eponate12 (Ted Pella). Sections were cut 60 nm thin onto formvar-filmed copper grids, stained with 2% uranyl acetate and Reynolds lead citrate, and imaged at 80 kV in a Zeiss EM10C (Oberkochen) equipped with an ES1000W Erlangshen CCD camera (Gatan).

### Genomic integration and FACS

HEK293-tetON and CHO-tetON cells were used for genomic integration of the mARGs. The cells were cultured in a 6-well plate containing 2 mL DMEM with 10% tetracycline-free FBS (Clonetech) and penicillin/streptomycin. Cells were transfected with the piggyBac transposon backbone containing the mARGs and the piggyBac transposase plasmid at a transposon:transposase molar ration of 2.5:1. Transfection was conducted using parameters mentioned above and the cells were allowed to incubate for 72 hours. Cells were induced with 1 µg/mL of doxycycline 24 hours prior to FACS (BD FACSAria III). Polyclonal subpopulations of mARG-expressing HEK293-tetON cells were sorted into the following four bins: (subtype 1) cells with eBFP2 fluorescence greater than 10^4^ and EmGFP fluorescence greater than 10^4^ and mCherry fluorescence greater than 2×10^4^ au, (subtype 2) cells with eBFP2 fluorescence between 3×10^3^ and 2×10^4^ and EmGFP fluorescence between 2×10^3^ and 2×10^4^ and mCherry fluorescence between 2×10^3^ and 2×10^4^ au, (subtype 3) cells with eBFP2 fluorescence between 10^3^ and 6×10^3^ and EmGFP fluorescence between 2×10^2^ and 10^3^ and mCherry fluorescence greater than 2×10^4^ au, (subtype 4) cells with eBFP2 fluorescence greater than 10^4^ and EmGFP fluorescence greater than 2×10^4^ and mCherry fluorescence between 2×10^3^ and 2×10^4^ au. CHO-tetON cells were transfected with mARGs and the piggyBac transposase plasmid similar to above. mARG-expressing CHO-tetON cells with eBFP2 fluorescence greater than 10^4^, EmGFP fluorescence greater than 10^4^ and mCherry fluorescence greater than 2×10^4^ au were sorted.

For monoclonal cell lines, naïve HEK293-tetON cells were transfected with mARGs and the piggyBac transposase similar to above. mARG-expressing cells with eBFP2 fluorescence greater than 10^4^, EmGFP fluorescence greater than 10^4^ and mCherry fluorescence greater than 2×10^4^ au were sorted. 576 cells were sorted in individual wells of 96-well plate and the surviving 30 cells were analyzed for gas vesicle expression as described above.

To generate mCherry-HEK cells, HEK293-tetON cells were transfected with piggyBac transposon plasmid containing TRE3G promoter driving mCherry and the transposase plasmid similar to above. mCherry-HEK cells were sorted from cells with mCherry fluorescence between 1.5×10^4^ and 10^5^ au.

### In vitro toxicity assays

The viability of the mARG-expressing cells was determined using three different assays involving cellular metabolic activity (resazurin reduction, MTT assay), quantification of cellular ATP content (CellTiter-Glo, Promega Corp.), and dye exclusion (Trypan Blue, Caisson Labs). The measurements were all quantified as percent viability compared to control cells that expressed mCherry only (mCherry-HEK). For the MTT and CellTiter-Glo assays, cells were grown in 96-well plates and induced with 1 µg/mL doxycycline and 5 mM sodium butyrate for 72 hours. They were then treated with reagents according the manufacturers’ protocols. Luminescence (CellTiter-Glo) and absorbance at 540 nm (MTT) was measured using a SpectraMax M5 spectrophotometer (Molecular Devices). For the Trypan Blue assay, the cells were first grown in 6-well plates and treated with 1 µg/mL doxycycline and 5 mM sodium butyrate for 72 hours. They were then trypsinized and resuspended in media before being stained 1:1 with Trypan Blue dye. Ten µL of the solution was loaded in a disposable hemocytometer (C-chip DHC S02, Incyto) and total cell count and blue-stained dead cells were quantified by bright field microscopy. Cellular morphology was imaged from mARG-HEK and mCherry-HEK cells after 3 days of expression with 1 µg/mL doxycycline and 5 mM sodium butyrate. Phase images were acquired using a Zeiss Axio Observer with a 20x objective.

### In vitro ultrasound imaging

To create phantoms for *in vitro* ultrasound imaging, wells were casted with molten 1% w/v agarose in PBS using a custom 3D-printed template. mARG-HEK and mCherry-HEK cells were allowed to express their transgenes using the specified inducer concentrations and expression duration. They were then trypsinized and counted via disposable hemocytometers in bright field microscopy. Next, cells were mixed at a 1:1 ratio with 50 °C agarose and loaded into the wells before solidification. The volume of each well was 60 µl and contained 6×10^6^ cells. The phantoms were submerged in PBS, and ultrasound images were acquired using a Verasonics Vantage programmable ultrasound scanning system and L22-14v 128-element linear array transducer with a 0.10-mm pitch, an 8-mm elevation focus, a 1.5-mm elevation aperture, and a center frequency of 18.5 MHz with 67% −6 dB bandwidth (Verasonics, Kirkland, WA). Each frame was formed from 89 focused beam ray lines, each with a 40-element aperture and 8 mm focus. A 3-half-cycle transmit waveform at 17.9 MHz was applied to each active array element. For each ray line, the amplitude modulation (AM) code was implemented using one transmit with all elements in the active aperture followed by 2 transmits in which first the odd- and then the even-numbered elements are silenced (*10*). Each image captured a circular cross-section of a well with a 4-mm diameter and center positioned at a depth of 8 mm. In AM mode, the signal was acquired at 0.27 MPa (2V) for 10 frames and the acoustic pressure was increased to 1.57 MPa (10V) to collect 46 additional frames. Ultrasound images were constructed by subtracting the collapsing frame by frame 4 post-collapse.

For Fig. 4, F-H, the high gas vesicle content of some samples resulted in acoustic shielding and a residual amount of gas vesicles remained intact after 46 frames of insonation at 1.57 MPa. To fully collapse all the gas vesicles and collect the background signal, the acoustic pressure was increased to 3.2 MPa (25V), then a second set of images was acquired with 10 frames at 0.27 MPa and 46 frames at 1.57 MPa. Gas vesicle-specific signal was determined by subtracting the total ultrasound signal from the 46 frames acquired before 3.2 MPa ultrasound by the total ultrasound signal from the 46 frames post collapse.

### Cytotoxicity assay on cells exposed to ultrasound

mARG-HEK and mCherry-HEK cells were cultured on custom made Mylar-bottom 24-well plates. Cells were cultured on fibronectin-coated Mylar films until they reached 80% confluency and induced for gas vesicle expression (1 µg/mL doxycycline and 5 mM sodium butyrate) for 3 days. The cells were then insonated from the bottom using an L22-14v 128-element linear array transducer (Verasonics). The transducer was mounted on a computer-controlled 3D translatable stage (Velmex). The bottom of the plates was acoustically coupled to the transducer with water and positioned 8 mm away from the transducer face. The cells were exposed to 3.2 MPa of pressure and the transducer was translated at a rate of 3.8 mm/s. The plates were returned to the incubator for 24 hours. Cytotoxicity was then assayed using resazurin reduction (MTT) on cells exposed to ultrasound and compared to non-insonated control cells.

### 3D cell culture and in vitro acoustic recovery after collapse

mARG-HEK and mCherry-HEK cells were mixed in Matrigel (Corning) containing 1 µg/mL of doxycycline and 5 mM sodium butyrate. The cell-laden hydrogels were placed in a 1% w/v agarose base to prevent cell migration out of the hydrogel and separate the cells away from the bottom of the plates during imaging. Cells were cultured for total of 6 days and imaged every 3 days from the top using an L22-14v 128-element linear array transducer (Verasonics). The transducer was wiped with 70% ethanol, and imaging was conducted in a laminar flow biosafety cabinet to preserve sterility. After imaging, to ensure complete collapse of all gas vesicles in the cells, the entire hydrogel was exposed to 3.2 MPa ultrasound and the transducer was translated three times across the gel at a rate of 1-2 mm/s. The culture media was changed daily and contained 1 µg/mL of doxycycline and 5 mM sodium butyrate.

### In vivo expression of gas vesicles and ultrasound imaging

All *in vivo* experiments were performed on NOD SCID mice (NOD.CD17 *Prkdc*^*scid*^/*NCrCr*l; Charles River), aged 10-15 weeks, under a protocol approved by the Institutional Animal Care and Use of Committee of the California Institute of Technology. mARG-HEK and mCherry-HEK cells were cultured in tetracycline-free media in T225 flasks. 1-1.2 x 10^7^ cells were trypsinized and the 200 µl cell-pellet was mixed with 200 µl Matrigel (Corning) containing 5 mM sodium butyrate. The mixture of mARG-HEK cells and Matrigel was injected subcutaneously in the left flank of mice and the mixture of mCherry-HEK cells and Matrigel was injected subcutaneously in the right flank of mice. Starting from the day of tumor inoculation, mice we interperitoneally injected with 200 µl of saline containing 75 µg doxycycline and 25 mg of sodium butyrate daily. The lower half of mice were depilated to allow for fluorescence imaging and ultrasound coupling.

For ultrasound imaging, the mice were anesthetized with 2% isoflurane and maintained at 37 °C using a heating pad. Ultrasound imaging was carried out using the pulse sequence described above with an L22-14v transducer attached to a custom-made manual translation stage. Using B-mode ultrasound imaging, the center of the tumor was positioned approximately 8 mm from the surface of the transducer, and gas vesicle-specific ultrasound images were acquired. The transducer was translated laterally with 1 mm steps to collect ultrasound images of most of the tumor.

High framerate ultrasound datasets for Doppler imaging were acquired with the same ultrasound transducer and scanner. The Doppler pulse sequence consisted of 11 tilted plane wave transmissions (varying from −10 to 10 degrees) at a 5.5 kHz framerate, leading to a 500 Hz framerate after coherent compounding. Plane wave transmissions lasted 0.5 s (or 250 frames). A power Doppler image representing blood flow was computed from each ensemble of 250 frames using a singular value decomposition filter that separates clutter from red blood cell echoes (*24*).

To obtain tissue samples after the mice were euthanized, tumors were resected and placed in 3.7% formaldehyde solution (4°C) for 24 hours and transferred to sterile 30% sucrose for an additional 24 hours. Tumors were embedded in OCT compound (Tissue-Tek), flash frozen and sectioned to 60 µm slices using a Cryostat (Leica CM3050). Sections were stained with TO-PRO3 nucleus stain, mounted (Fluoromount Aqueous Mounting Medium) and imaged using a Zeiss LSM 800 confocal microscope.

## ACKNOWLEDGEMENTS

The authors thank David Maresca, Bill Ling and Avinoam Bar-Zion for help with ultrasound imaging, Noushin Koulena for assistance with tissue histology, Andres Collazo for confocal microscopy, Chris Buser and Oak Crest Institute of Science for cell sectioning and staining, Erik Jue and William Chour for assistance with initial experiments. Electron microscopy was performed at the Beckman Institute Resource Center for Transmission Electron Microscopy at Caltech. Fluorescence imaging of tissues was performed in the Biological Imaging Facility of the Caltech Beckman Institute with support from the Arnold and Mabel Beckman Foundation. We appreciate the help of Audrey Lee-Gosselin and Caltech’s Office of Laboratory Animal Research with animal protocols and husbandry. Arash Farhadi was supported by the NSERC graduate fellowship. Daniel Sawyer was supported by the NSF graduate research fellowship (Grant No. 1745301). This research was supported by the National Institutes of Health (grant numbers R01EB018975 and U54CA199090 to M.G.S.), the Heritage Medical Research Institute (M.G.S.), the Packard Fellowship for Science and Engineering (M.G.S.), and the Burroughs Welcome Fund Career Award at the Scientific Interface (M.G.S.).

## AUTHOR CONTRIBUTIONS

A.F. and M.G.S. conceived and planned the research. A.F. and G.H.H. performed the experiments. A.F. and R.W.B. designed the DNA sequences. A.F. and D.P.S. designed and optimized the ultrasound imaging sequences. A.F. analyzed data. A.F., and M.G.S. wrote the manuscript with input from all authors. M.G.S. supervised the research.

