## Supplementary Figures and Tables for "Ultrasound Imaging of Gene Expression in Mammalian Cells"

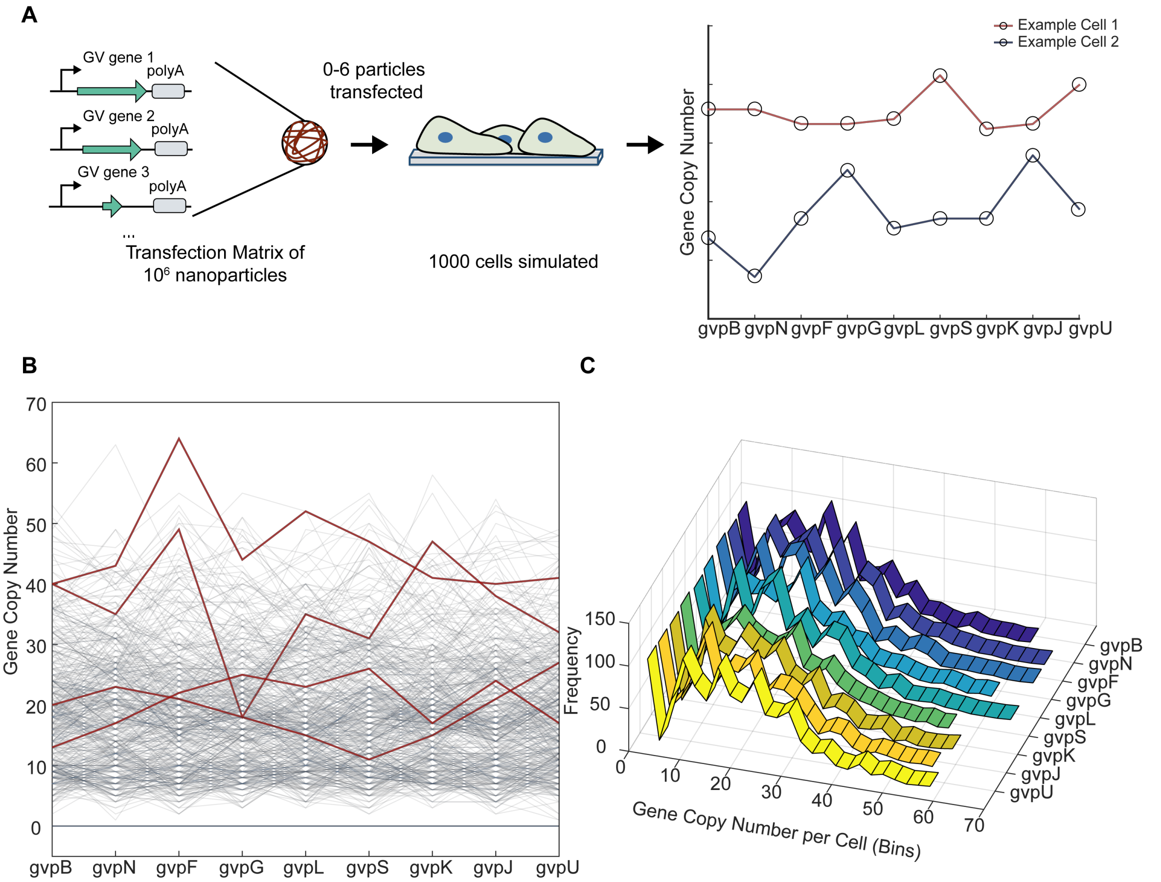


**Supplementary Figure 1 – Simulation of stochastic transfection assay.** (**A**) Schematic of the transfection process being simulated. 9 unique genes are randomly complexed with equal probability into polyethyleneimine-DNA transfection particles, each containing ~90 plasmids. The plasmids are then applied to cells, with each cell receiving 6 transfection particles, each particle with a 33% probability of contributing to gene expression. Values for this simulation were estimated from Bhise et al. (*1*) and Cohen et al. (*2*). (**B**) Pattern of gene copy number transfected to cells, with each gray line representing an individual cell. The red lines highlight 4 randomly chosen cells. (**C**) Histogram of the copy number of each gene transfected across 1000 cells. These results illustrate the stochasticity of transient multi-plasmid transfection and are not meant to serve as precise estimates of plasmid transfection in our experiments.


bioRxiv preprint. March 18, 2019. All rights reserved.

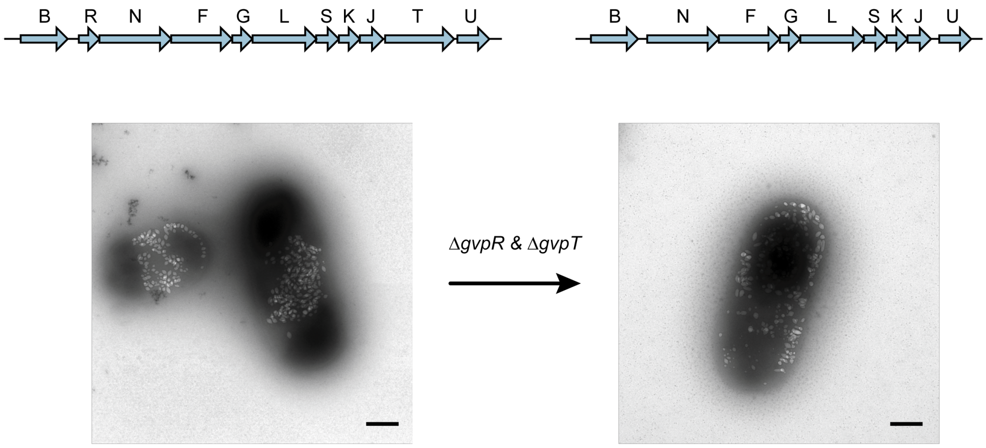


**Supplementary Figure 2 – *gvpR* and *gvpT* genes in the *B. megaterium* gene cluster are not necessary for gas vesicle formation.** Schematic of bacterial gas vesicle gene clusters used for heterologous expression of gas vesicles in *E. coli* (top). Representative whole cell TEM images of *E. coli* Rosetta 2(DE3)pLysS cells after expression of gas vesicles genes for 22 hours (bottom). Scale bars represent 500 nm. Expression performed as in Farhadi *et al.* (*3*) and TEM imaging as in Bourdeau et al (*4*).


bioRxiv preprint. March 18, 2019. All rights reserved.

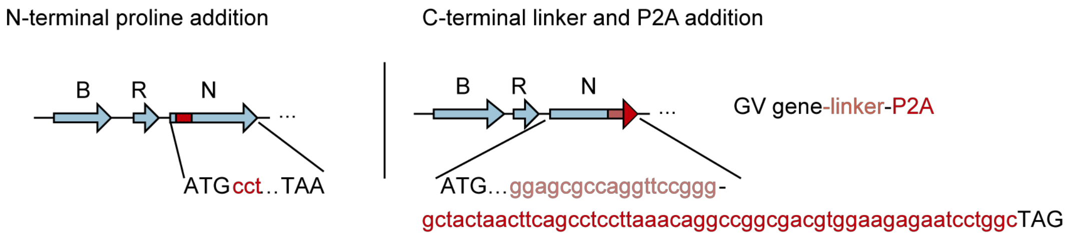


**Supplementary Figure 3 – Assay for tolerability of P2A peptide additions.** Illustration of gas vesicle gene cluster with N- or C-terminal modifications of each gene to test tolerability of P2A peptides, tested one-by-one in *E. coli*.

| Gene | GVs after N-term addition? | GVs after C-term addition? |
|---|---|---|
| *gvpB* | -- | No |
| *gvpR* | Yes | Yes |
| *gvpN* | Yes | Yes |
| *gvpF* | Yes | Yes |
| *gvpG* | Yes | Yes |
| *gvpL* | Yes | Yes |
| *gvpS* | Yes | Yes |
| *gvpK* | Yes | Yes |
| *gvpJ* | Yes | Yes |
| *gvpT* | Yes | Yes |
| *gvpU* | Yes | Yes |

**Supplementary Table 1 – Tolerability of P2A peptide additions to *B. megaterium* gas vesicle genes.** Each gene of the *B. megaterium* gene cluster was modified with an N-terminal proline after the start codon or with a linker and P2A peptide at the C-terminus, resulting in a total of 21 unique GV gene clusters. *E. coli* were transformed with each plasmid and gas vesicles were induced for expression for a total of 22 hours and assayed for the presence of gas vesicles using TEM. The table indicates whether gas vesicles were observed by TEM. Expression and TEM imaging performed as in Farhadi *et al.* (*3*).


bioRxiv preprint. March 18, 2019. All rights reserved.

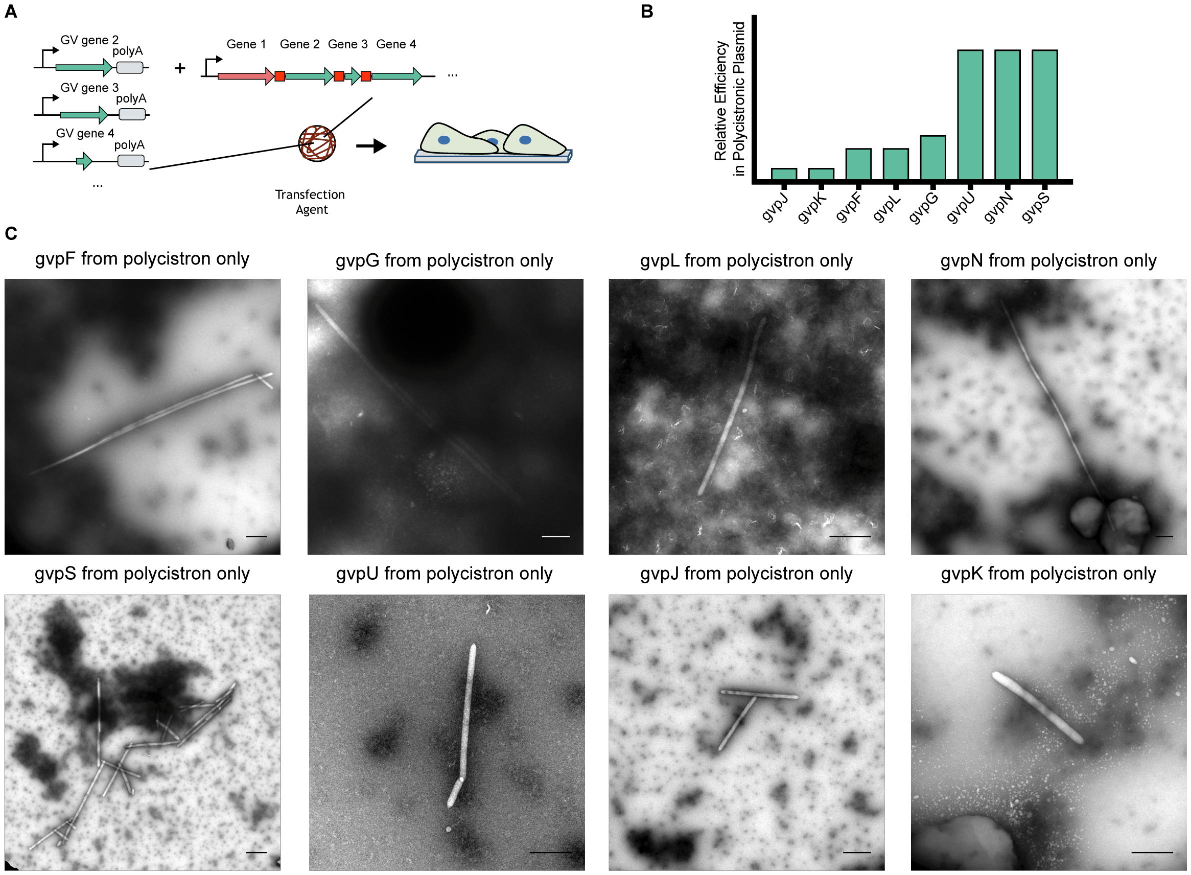


**Supplementary Figure 4 – Identification of bottleneck genes on the polycistronic gas vesicle gene plasmid.** (**A**) Schamatic of the experiment. To test the efficiency with which gas vesicles could be formed when a given gene was supplied only on the polycistronic plasmid, cells were co-transfected with a monocistronic plasmid containing GvpB, 7 other monocistronic plasmids including all but the gene being assayed, and the polycistronic plasmid. (**B**) Qualitative estimate of the relative number of gas vesicles produced when each indicated gene was supplied solely by the polycistrinic plasmid. (**C**) Representative TEM images of gas vesicles in the lysate of HEK293T cells for all 8 assays. Scale bars represent 500 nm.


bioRxiv preprint. March 18, 2019. All rights reserved.

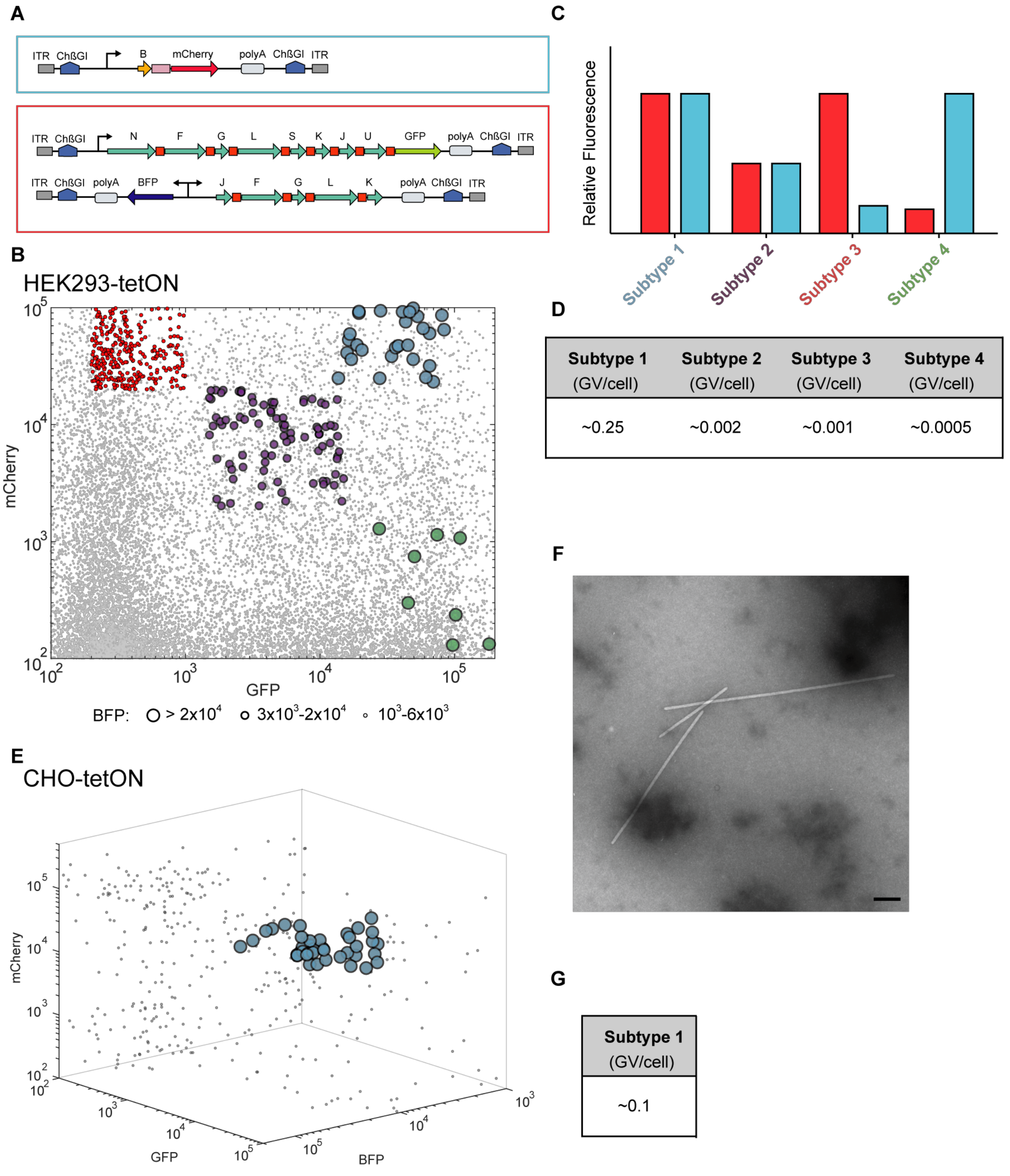


| Subtype 1 (GV/cell) | Subtype 2 (GV/cell) | Subtype 3 (GV/cell) | Subtype 4 (GV/cell) |
|---|---|---|---|
| ~0.25 | ~0.002 | ~0.001 | ~0.0005 |

| Subtype 1 (GV/cell) |
|---|
| ~0.1 |

**Supplementary Figure 5 – Fluorescence activated cell sorting of HEK293-tetON and CHO-tetON cells transfected with integrating mARG contructs.** (**A**) Diagram of the integrating constructs used to generate polyclonal cell lines. (**B**) FACS of mARG-expressing HEK293-tetON cells. Colored data indicate cells sorted for each group and gray dots are unsorted population. (**C**) Illustration of the four polyclonal subtypes sorted to study the impact of polycistron stoichiometry on gas vesicle expression. Red bars indicate mCherry expression; cyan bars indicate EmGFP and eBFP2 expression. (**D**) Approximate gas vesicle yield from polyclonal cells in each subtype. (**E**) FACS of mARG-expressing CHO-tetON cells. Colored data indicate cells sorted in subtye 1 and gray dots are unsorted cells. (**F**) Representative TEM image of bouyancy-enriched lysate from CHO-tetON cells sorted as indicated in (E). Scale bar represeents 500 nm. (**G**) Approximate gas vesicle yield for the sorted mARG-expressing CHO-tetON cells.


bioRxiv preprint. March 18, 2019. All rights reserved.

| Collected from FACS | Formed colonies | Triple positive fluorescence | Formed GVs (TEM) | >1 GVs/cell |
|---|---|---|---|---|
| 576 | 30 | 21 | 12 | 6 |

**Supplementary Table 2 – Selection funnel for monoclonal mARG-HEK cells.** The numbers indicate the number of cells or cell lines selected at each stage.

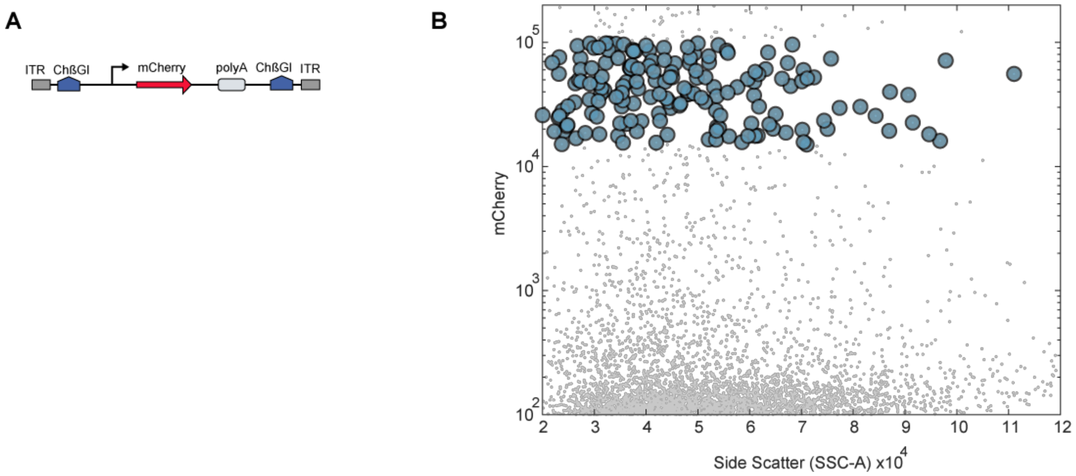


**Supplementary Figure 6 – Genetic construct and sorting of mCherry-HEK cell line.** (**A**) Genetic construct for stable genomic integration of mCherry containing a TRE3G promoter upstream and SV40 polyadenylation element downstream of mCherry. (**B**) FACS of mCherry cells, with selected cells indicated with blue dots.

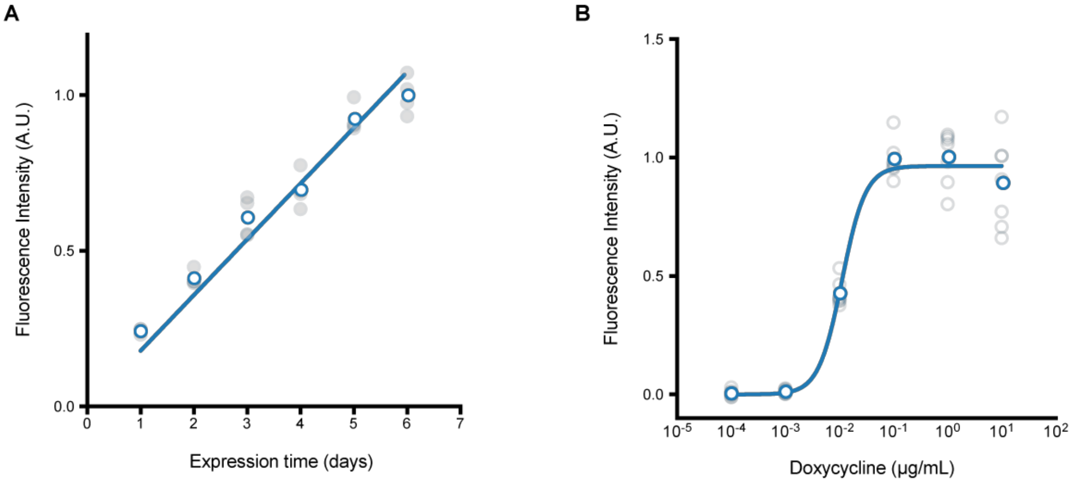


**Supplementary Figure 7 - Fluorescence measurements of gene expression as a function of time and inducer concentration in mARG-HEK cells.** (**A**) mCherry fluorescence of mARG-HEK cells induced with 1 µg/mL doxycycline and 5 mM sodium butyrate at the indicated times after induction (n=4, with the darker dots showing the mean). (**B**) mCherry fluorescence of mARG-HEK cells with the indicated inducer concentration and 5 mM sodium butyrate after 72 hours of induction (n=7, with the darker dots showing the mean).



bioRxiv preprint. March 18, 2019. All rights reserved.

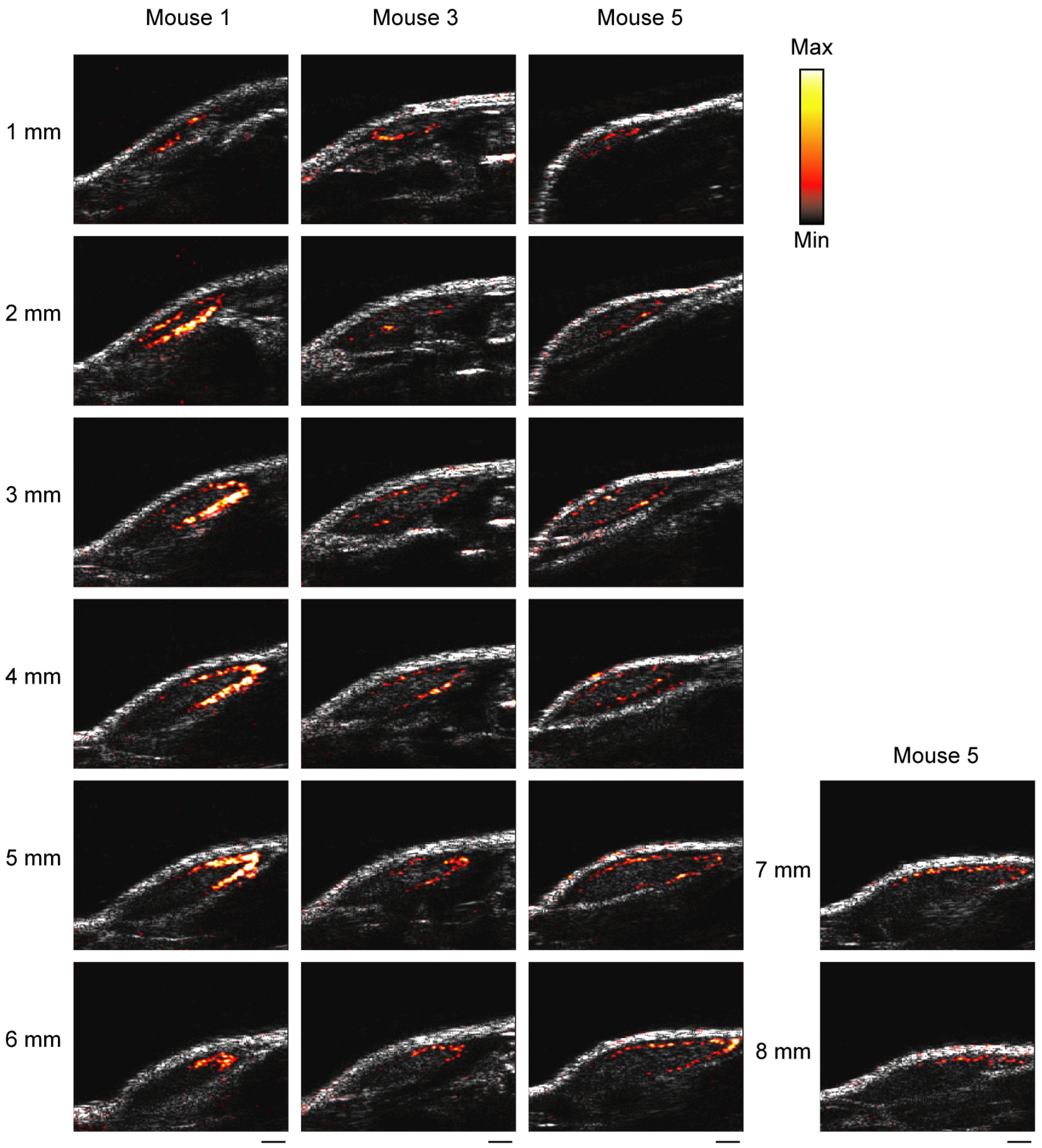


**Supplementary Figure 8 - Additional examples of *in vivo* ultrasound images of adjacent planes in mARG-HEK tumors acquired at 1 mm intervals.** For each imaging slice the difference heatmap of nonlinear signal between frame 1 and frame 4 is overlaid on grayscale anatomical scale. Minimum and maximum values of colorbar are 4000 and 40000, respectively. Scale bars are 1 mm.



bioRxiv preprint. March 18, 2019. All rights reserved.

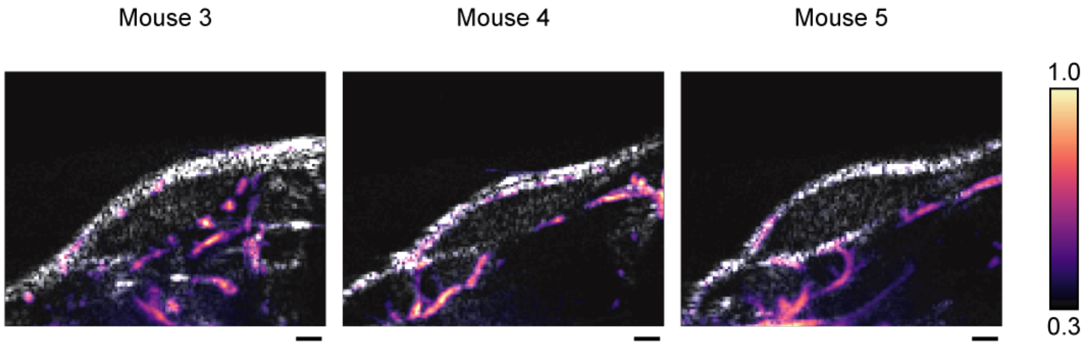


**Supplementary Figure 9 – Representative Doppler images of tumors containing mARG-HEK cells.** Doppler ultrasound images were acquired using 250 frames of ultrafast planewaves at 25V (3.2 MPa) and used to reconstruct vascular maps ploted as normalized power doppler signal overlaid on anatomical images in grayscale. Scale bars represent 1 mm.

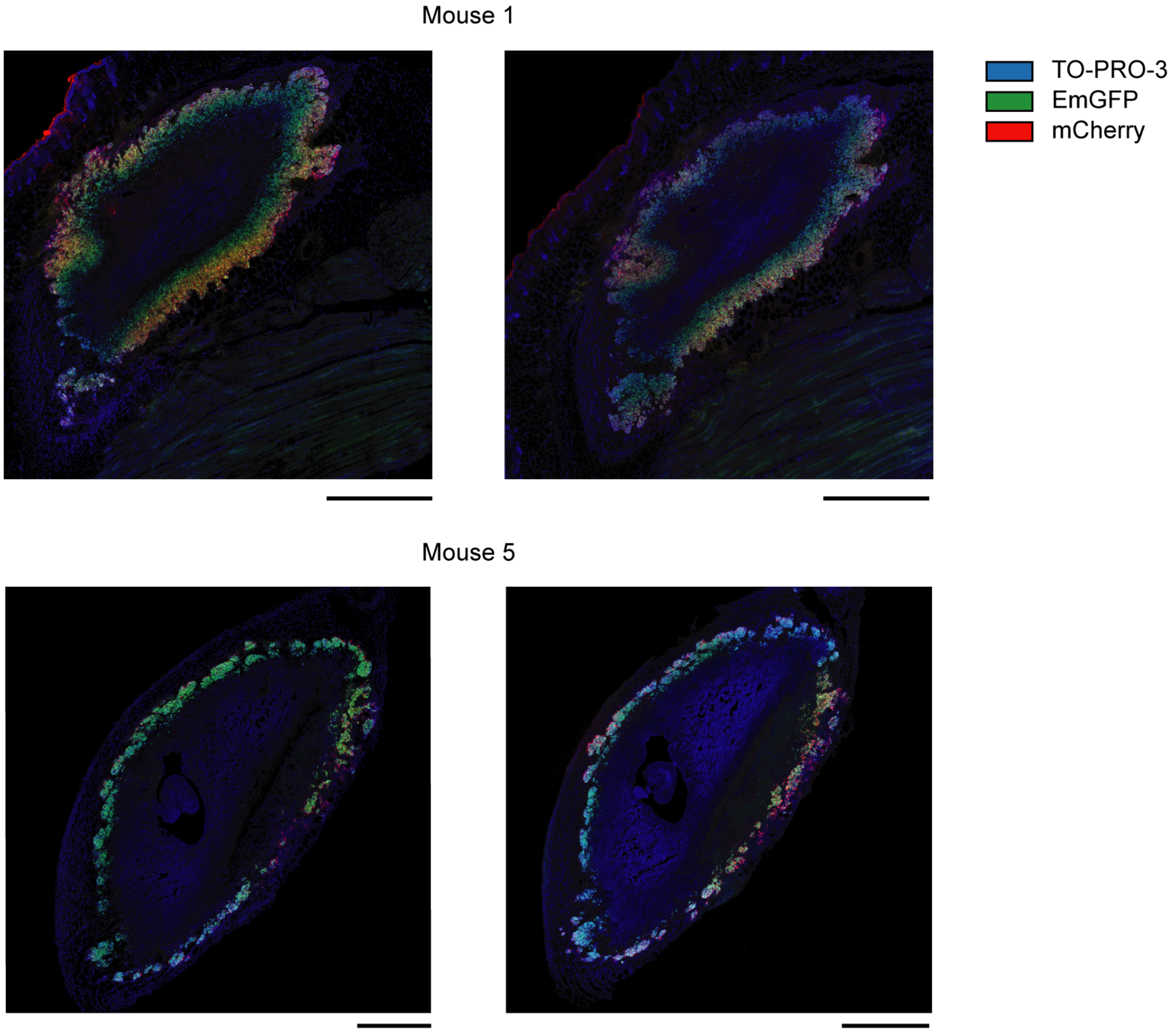


**Supplementary Figure 10 – Representative histology sections of tumors containing mARG-HEK cells.** For each mouse, two neighboring sections are presented. Blue color indicates cell nuclear staining using TO-PRO3, green color represents GFP fluorescence and red color represents mCherry fluorescence. All scale bars are 1 mm.


bioRxiv preprint. March 18, 2019. All rights reserved.

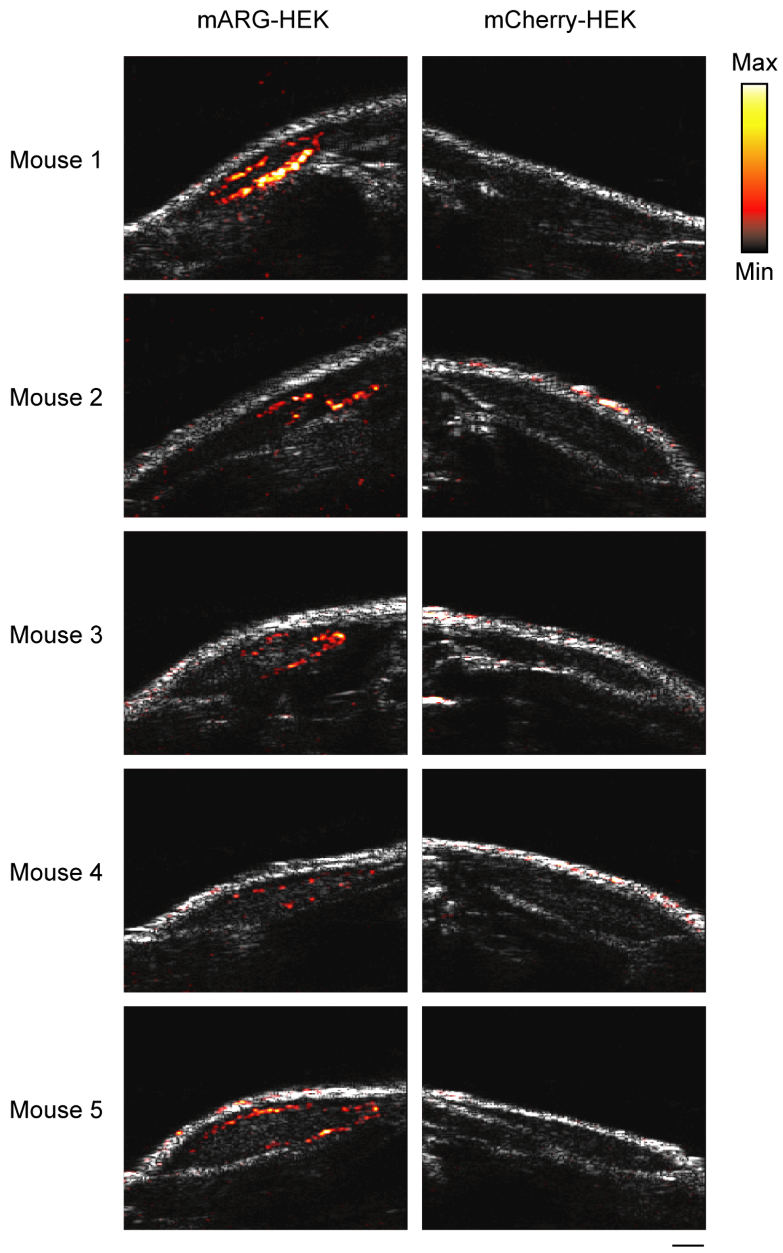


**Supplementary Figure 11 – Biological replicates of *in vivo* ultrasound imaging of gene expression.** The left column shows ultrasound images of tumors containing mARG-HEK cells after 4 days of doxycycline administration. The right column shows ultrasound images of tumors containing mCherry-HEK cells after 4 days of doxycycline administration. Difference heatmap of nonlinear signal between frame 1 and frame 4 is overlaid on a grayscale anatomical ultrasound image. Min and max on colorbar represent 4000 and 40000, respectively. The scale bar represents 1 mm.


bioRxiv preprint. March 18, 2019. All rights reserved.

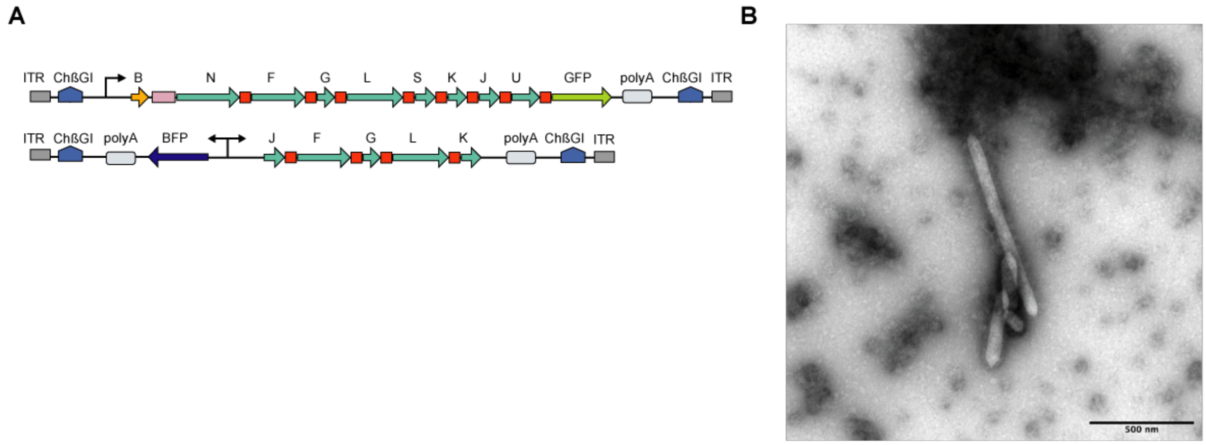


**Supplementary Figure 12 – Consolidated mARG construct comprising 2 gene cassettes enables mammalian gas vesicle expression.** (**A**) Schematic of two gene cassettes integrated to the genome of HEK293-tetON cells. In the top construct *gvpB* is separated from *gvpN* by an internal ribosome entry sequence (shown in purple). (**B**) Representative TEM image of GVs in the lysate of HEK293-tetON cells transfected with the constructs in (A) and induced with 1 μg/mL doxycycline.


bioRxiv preprint. March 18, 2019. All rights reserved.


bioRxiv preprint. March 18, 2019. All rights reserved.
